# Disrupted Brain Organoid Circuitry, Structural Organization, and Spine Morphology in 7q11.23 Copy Number Variant Syndromes

**DOI:** 10.64898/2026.08.11.744287

**Authors:** Inyoung Hwang, Jasmine S. Yeo, Maria Camila Almeida, Bárbara Mejía Cupajita, Daniel Carneiro Carrettiero, Juliana Acosta-Uribe, Albert Han, Anna Ngo, Carolina Camargo, Magdalena Budisteanu, Aurora Arghir, Lucy R Osborne, James Ellis, Ikuko T. Smith, Michael J. Goard, Kenneth S. Kosik

## Abstract

The 7q11.23 chromosomal region represents a model of gene dosage-dependence, where a hemizygous deletion causes Williams Syndrome (WS) and a duplication leads to 7q11.23 Duplication Syndrome (Dup7). It is not understood how these copy number variations (CNVs) disrupt development and functional cortical circuit assembly. Utilizing iPSC-derived cerebral organoids and longitudinal imaging from post-differentiation day 30 to 150, we characterized the aberrant neural rosette morphogenesis in WS and Dup7 during early stages, establishing an early structural divergence from control lines. We observed accelerated early cortical rosette morphogenesis in WS, characterized by a premature increase in both rosette number and layer thickness compared to controls. In contrast, Dup7 organoids consistently exhibit a significantly lower number and reduced thickness of rosettes from early stages onward. This early structural disruption progressively impacted synaptic-level architecture as the organoids matured. Dendritic spine characterization at later stages revealed Dup7 organoids exhibited a significantly higher dendritic spine density compared to WS. Pharmacological antagonism of CCR5 (C-C chemokine receptor type 5) with Maraviroc significantly enhanced dendritic spine density in control and WS organoids; however, this effect was absent in Dup7. To determine how these structural anomalies translate into circuit-level behavior, we performed longitudinal calcium imaging using GCaMP. Control organoids sustained synchronized activity and high spike correlations at all time points. This synchronization was delayed and highly transient in WS organoids, and completely abolished in Dup7 organoids, which exhibit significantly low spike correlations at all stages. Developmentally, GABA changes from acting as an excitatory signal in the immature brain to an inhibitory signal as the brain matures. As control lines matured gabazine-induced desynchronization progressively diminished, but persisted in WS organoids. Dup7 organoids failed to establish synchronization at any developmental time point, but displayed a negligible increased synchrony following gabazine treatment. These functional aberrations were paralleled by genotype-specific defects in structural organization, specifically in rosette morphogenesis and dendritic spine density. Collectively, our findings demonstrate that 7q11.23 CNVs trigger pathogenic neurodevelopmental defects by derailing the trajectories of structural organization, circuit assembly, and functional synchronization during cortical maturation.

## Introduction

Williams syndrome (WS) and 7q11.23 duplication syndrome (Dup7) are reciprocal copy number variation (CNV) disorders caused by heterozygous deletion or duplication, respectively, of a ∼1.5 Mb genomic region containing approximately 28 genes^1–12^. Altered gene dosage at this locus produces contrasting behavioral and neurodevelopmental phenotypes: WS is associated with hyper-sociability, relative verbal strengths, and microcephaly, whereas Dup7 often presents with social withdrawal, speech delay, and macrocephaly^1–18^. Previous investigations have interrogated the influence of individual genes within this genomic region, establishing that single-gene manipulations can independently influence alterations in neural differentiation dynamics, various molecular pathways and behavioral phenotypes^3–5,8–10,13–16,19–24^. While these studies have offered gene-specific insights, a comprehensive characterization of how these divergent genetic influences collectively disrupt the temporal coordination of structural and functional development remains a critical gap in our understanding.

Human induced pluripotent stem cell (iPSC)-derived cerebral organoids provide a patient-specific platform for interrogating both the structural and functional dynamics of cortical development^25–27^. This system recapitulates features of early neurodevelopmental milestones *in vitro* by inducing the formation of neuroepithelial rosettes—radially organized progenitor structures that reflect early cortical patterning and ventricular zone–like organization^25^. Although isolated reports suggest mature dendritic spines can be observed beyond day 150, we lack a comprehensive characterization of spine morphology across these genotypes^32^. Maraviroc, a selective antagonist of CCR5 (C-C chemokine receptor type 5) is known to promote dendritic spine maturation^31,34–36^ and was shown here to have genotypic effects.

We proceeded from an anatomical analysis of the WS, Dup7, and control genotypes to a physiological circuitry analysis using GCaMP imaging by 2-photon microscopy. The spontaneous emergence of highly pre-configured activity in human cerebral organoids^37^ in which the participating units are highly correlated as measured on multi-electrode arrays^36^ suggest that they can offer insights into the pathophysiology of neurodevelopmental disorders. Organoid development is situated at a key developmental transition in GABAergic signaling with some parallels to early mammalian cortical development. Episodic synchronous network events coordinate neuronal assemblies and pattern the neocortex into functionally related modules^38–41^. These early synchronous oscillations are largely driven by a depolarizing GABAergic program^42–44^, as immature GABAergic signaling provides excitatory input that organizes large-scale network bursts^40,42,44–47^. As cortical circuits mature, GABAergic signaling transitions from depolarizing to hyperpolarizing, and inhibitory interneuron networks expand^39,40,42,44,45,45,48^. This developmental shift promotes desynchronization of neuronal firing, enabling more efficient and flexible information processing^40,44,45,49^. The precise timing of this transition—from GABA-driven synchrony to inhibition-supported desynchrony—is critical for establishing functional cortical circuitry, and disruptions to this sequence are increasingly implicated in neurodevelopmental disorders^38–40,44,50–52^. In this study, we define the coordinated trajectories of network activity, rosette organization, and spine maturation, elucidating how 7q11.23 gene dosage affects the expected synchronization of neural activity. Our findings revealed that 7q11.23 dosage imbalance disrupts neurodevelopmental progression by decoupling early structural organization from late-stage functional synchronization.

## Methods

### Generation and Characterization of iPSCs

Control #1 iPSC line (F12442.4) was donated by Karch^53^. Control #2 iPSC line (KOLF2.1J) was purchased from The Jackson Laboratory. WS1 and WS2 were obtained from skin biopsies from two females at the Alexandru Obregia Clinical Hospital of Psychiatry in Bucharest, Romania and Victor Babes National Institute of Pathology^22^. WS1 and WS2 iPSC lines were reprogrammed from skin fibroblasts obtained from skin biopsies collected following written informed consent from the donors and approved by the Washington University School of Medicine Institutional Review Board and Ethics Committee (IRB 201104178, 201306108). WS3 fibroblasts were obtained from the SickKids Heart Centre Biobank and reprogrammed into iPSCs using retroviral vectors including the EOS pluripotency reporter vector^54^, with an initial characterization^55^. Dup1 and Dup2 iPSC lines were provided from Osborne Lab of University of Toronto, Canada. For all 7q11.23 CNV cell lines used in this study, we performed Next-Generation Sequencing (NGS) to rigorously identify and confirm the respective deletions and duplications. Detailed genomic identification data for these lines can be found in Supplementary Table 1. Briefly, All the iPSC lines were cultured in mTeSR1 medium (Stem Cell Technologies, # 100-0276) on tissue culture plates coated with hESC-qualified Matrigel (Corning, # 354230). mTeSR1 was exchanged every other day and iPSCs were routinely passed using ReLeSR (Stem Cell Technologies, # 100-0483). To characterize each iPSC line, genomic analysis was commissioned to the Human Genome Sequencing Center (HGSC) at Baylor College of Medicine. Specifically, following sample quality control (QC), PCR-free short-read whole-genome sequencing libraries were prepared using KAPA Hyper PCR-free reagents (Roche, # KK8505). Final library size estimation and quantification were verified using a Fragment Analyzer electrophoresis system (Agilent Advanced Analytical Technologies, Inc.) and a QuantStudio 6 Flex Real-Time PCR System (Applied Biosystems), respectively. Libraries were then sequenced on an Illumina NovaSeq X platform to generate 150 bp paired-end reads in a multiplexed pool format, achieving an average sequencing depth of 10X. This approach was selected to minimize amplification bias and ensure high-fidelity detection of structural variants within the 7q11.23 locus (chr7:72,700,000–74,200,000; GRCh38). Post-sequencing data processing was performed via the HGSC research DRAGEN analysis pipeline. Primary base calling was executed using DRAGEN’s BCL Conversion functionality, and secondary analyses—including alignment to the GRCh38 human reference genome, indel variant calling, and coverage QC metric generation—were completed using the DRAGEN platform (version 4.3.6) configured for DNA Germline WGS. The gender and age of each donor, along with the precise 7q11.23 deletion or duplication boundaries for each iPSC line, are provided in Supplementary Table 1, and raw datasets are provided in Data S1.

### Cerebral Organoid Generation

Cerebral organoids were generated using a modified version of the protocol from Pasca Lab^25,56,57^. Human induced pluripotent stem cells (hiPSCs) were enzymatically dissociated using Accutase (Gibco, A1110501) and seeded at a density of 10,000 cells per well into 96-well slit-well plates (S-Bio, MS-9096SZ) in mTeSR™1 medium (STEMCELL Technologies, 100-0276) supplemented with 1:1000 ROCK inhibitor (Tocris, 1254/10). After overnight incubation, the medium was changed daily from day 1 to 5 with Media A, composed of E6 medium (Gibco, A1516401), 10 µM SB431542 (Tocris, 1614), 2.5 µM Dorsomorphin (Tocris, 3093), 2.5 µM XAV-939 (Tocris, 3748), and 1x Anti-Anti (Gibco, 15240062). From day 6 to 15, cultures were maintained in Media B with daily media changes, consisting of Neurobasal-A medium (Gibco, 12349015), B-27 Supplement minus vitamin A (Gibco, 12587001), GlutaMAX (Gibco, 35050061), Anti-Anti (Gibco, 15240062), 20 ng/ml bFGF (Shenandoah Biotechnology, 100-28), and 20 ng/ml EGF (Shenandoah Biotechnology, 100-26). From day 16 to 24, organoids were maintained in the same Media B, with media changes every other day. Beginning on day 20, half of the organoids were transferred to anti-adhesive-treated 6-well plates. From day 25 to 42, organoids were cultured in Media C, which included Neurobasal-A, B-27 without vitamin A, GlutaMAX, Anti-Anti, and 20 ng/ml each of NT-3 (Shenandoah Biotechnology, 800-06) and BDNF (Shenandoah Biotechnology, 100-01), with media changes every other day. After day 43, organoids were maintained in Media D, which had the same basal composition but without growth factors, and was refreshed every 3–4 days.

For CCR5 antagonism studies, Maraviroc^31,35,36,58^ was administered to the treatment group. Starting at Day 120, organoids were treated with 200 ng/mL of Maraviroc every other day for a total duration of 30 days^58^.

### Whole-mount and Sectioned Organoids for Immunohistochemistry

Organoids were processed for whole-mount immunostaining using the following reagents: 2% PBST (2% Triton X-100 in 1X PBS with 0.05% sodium azide), 1X PBS, blocking buffer (10% normal goat serum, 1% Triton X-100, 2.5% DMSO, 0.05% sodium azide in 1X PBS), antibody dilution buffer (1% normal goat serum, 0.2% Triton X-100, 2.5% DMSO, 0.05% sodium azide in 1X PBS), washing buffer (3% NaCl, 0.2% Triton X-100 in 1X PBS), primary and secondary antibodies, RapiClear® 1.52 (Sunjin Lab, RC152001), and iSpacer® chambers (Sunjin Lab, IS013). Organoids were fixed in 4% paraformaldehyde and incubated in 2% PBST on an orbital shaker for 2–3 days at 35–37 °C, followed by an additional 2–3 days of permeabilization in 2% PBST under the same conditions. Samples were washed in 1X PBS three times for 15 min at room temperature and then incubated in freshly prepared blocking buffer for 1–2 days at 4 °C. Primary antibodies were applied in antibody dilution buffer for 3–5 days at 4 °C. After incubation, samples were washed three times for 1 h in washing buffer at room temperature and then maintained in washing buffer overnight at 4 °C on an orbital shaker. Organoids were subsequently incubated with secondary antibodies in antibody dilution buffer for 2–3 days at 4 °C or room temperature, washed three times for 1 h in washing buffer, and placed overnight at 4 °C. Samples were then washed in 1X PBS three times for 20 min. For nuclear counterstaining, organoids were incubated overnight at 4 °C in Hoechst 33342 (Abcam, Ab228551) in 1X PBS (1:2500) and washed in 1X PBS three times for 1 hour at room temperature. Tissue clearing was performed with RapiClear® at room temperature for 1 day (pre-warmed to 37 °C to facilitate penetration), using a minimum of five times the sample volume. Cleared specimens were mounted in iSpacer® microchambers filled with fresh RapiClear®, and excess reagent was removed from edges with lint-free wipes.

Immunohistochemistry on cryosectioned or paraffin-embedded, microtome-sectioned organoids using a panel of primary antibodies against key developmental and regional markers. Organoids were first fixed in 4% paraformaldehyde (PFA) overnight at 4 °C on a shaker. Following fixation, they were cryoprotected in 30% sucrose in 1X PBS until fully equilibrated, then embedded and cryosectioned. Tissue sections were permeabilized in PBST (0.3% Triton X-100 in 1X PBS) for 1 hour at room temperature, followed by a 5-minute PBST wash on a room temperature shaker. Sections were then incubated in a blocking solution containing 10% normal goat serum (NGS) and 1% bovine serum albumin (BSA) in PBST to prevent nonspecific binding. Primary antibodies were applied in blocking solution and incubated overnight at 4 °C. Primary antibodies were applied at validated working concentrations and dilutions as follows: SATB2 (Rabbit pAb, 1:400; Abcam, ab34735), MAP2 (Rabbit pAb, 1:500, Abcam, ab32454; Mouse mAb, 1:1000, Thermo Fisher, 13-1500; Guinea Pig pAb, 1:1000, Synaptic Systems, 188004), CTIP2 (Rat mAb, 1:500; Abcam, ab18465), GFAP (Chicken pAb, 1:1000, Abcam, ab46741; Rabbit pAb, 1:1000, Abcam, ab7260), SOX2 (Mouse mAb, 1:200, Thermo Fisher, MA1-0144; Rabbit pAb, 1:500, Millipore, AB5603), SMI312 (Mouse mAb, 1:700; BioLegend, 837904), and TBR1 (Rabbit pAb, 1:500; Abcam, ab31940). Nuclei were counterstained using Hoechst 33342 (1:2500; Abcam, Ab228551). For visualization, species-specific Alexa Fluor™- conjugated secondary antibodies (Thermo Fisher Scientific) were applied at a 1:1000 dilution: Goat anti-Rabbit IgG Alexa Fluor™ 488 (A-11008), Goat anti-Guinea Pig IgG Alexa Fluor™ 594 (A-11076), Goat anti-Rat IgG Alexa Fluor™ 647 (A-21247), Goat anti-Mouse IgG Alexa Fluor™ 546 (A-11003), and Goat anti-Chicken IgG Alexa Fluor™ 647 (A-21449). Imaging was performed using a Leica SP8 resonant scanning confocal at 40X objectives and processed in ImageJ (Supplementary Fig. 4).

### Quantitative Anatomical Analysis of Organoid Features

To characterize the anatomical metrics of the organoids, we utilized two distinct quantification methods tailored to different developmental stages and structural scales. The first method was designed to track overall growth rates longitudinally during the early phase of rapid volume expansion (Day1 to Day55). For this, two-dimensional cross-sectional area of each live organoid was captured directly within the culture plate at 5-day intervals using a bright-field microscope. The total area was quantified by manually outlining the irregular outer boundary of the intact live organoid body using the Freehand Selection tool in ImageJ, where the built-in Measure Function computed the exact area based on the pre-set pixel.

The second method was employed to assess long-term changes in overall size alongside detailed internal anatomical features at specific fixed time points from day 30 through day 150 at 30-day intervals. Imaging was performed using a Leica SP8 resonant-scanning confocal microscope equipped with a 10X objective on fixed, tissue cleared, and immunohistochemistry (IHC)-stained whole mount organoids. Due to the large size of the organoids, multiple adjacent fields of view were acquired and automatically stitched together to capture both the entire tissue area for total size assessment and the internal anatomy. Image processing and quantification were conducted in Imaris. To determine the size of organoids, the diameter of the individual organoid was measured and subsequently converted into two-dimensional projected cross-sectional areas (A) by applying the area formula A= π x (diameter/2)^2^. For internal anatomical analysis, individual rosettes were identified within the 3D images. We measured the diameter of each rosette and its corresponding internal ventricular lumen. The average thickness of rosettes was then calculated by subtracting the lumen diameter from rosette diameter and dividing the remaining value by 2. These calculated values for rosette number and rosette thickness were then exported for statistical analyses in Prism 10 to compare metrics across genotypes at individual time points.

### AAV Transduction for GCaMP and GFP

Adeno-associated viral (AAV) vectors encoding GCaMP6m (Addgene, 100841-AAV9) and GFP (Addgene, 105558-AAV9; 51502-AAV9) were obtained from Addgene. Viruses were diluted to 0.2 × 10^11^ GC/mL to 0.2 × 10^12^ GC/mL in cerebral organoid growth medium immediately before use. Organoids retrieved two weeks prior to each imaging session were incubated overnight at 37 °C with 5% CO_₂_ in AAV-containing medium. The following day, organoids were subjected to a second overnight transduction in medium for optimal visualization with final titer between 1 × 10^10^ GC/mL and 1 × 10^11^ GC/mL, after which fresh medium was added before returning the cultures to standard conditions until imaging.

For GFP labeling in longitudinal dendritic spine imaging, organoids were incubated overnight at 37 °C with 5% CO_₂_ in medium containing AAV9-CAG-FLEX-EGFP-WPRE with final titer between 0.2 × 10^11^ GC/mL to 0.2 × 10^12^ GC/mL, together with AAV9-CamKII-0.4-Cre-SV40 with final titer between 0.1 × 10^10^ GC/mL to 0.5 × 10^11^ GC/mL.

### 2P Imaging for GCaMP Activity and Dendritic Spine Morphology

For two-photon (2P) live imaging, cerebral organoids were plated onto six-well plates pre-coated with poly-D-lysine (Gibco, Cat, A3890401) and 0.1% polyethylenimine. Following coating, plates were rinsed six times with ultrapure distilled water before organoid deposition. One day before each imaging session, organoids were transitioned to a 1:1 mixture of BrainPhys medium (STEMCELL Technologies, 05796) and cerebral organoid growth medium to optimize physiological firing conditions.

All imaging sessions were conducted within ±3 days of the designated developmental time points. GCaMP6m and GFP fluorescence were acquired using a Bruker Ultima 2P Plus microscope equipped with a resonant galvo-scanning module and controlled via PrairieView software. Fluorescence excitation was provided by an MKS Spectra-Physics InSight Dual S+A laser tuned to 920 nm, utilizing integrated DeepSee technology for automated dispersion compensation. Laser power at the sample was maintained between 60-225 mW(rarely and briefly at the high to capture spines) and optimized for GCaMP6m expression levels while keeping photobleaching below 1% min^-1^. To further minimize phototoxicity, laser exposure was strictly regulated by a High-Speed Shutter Control unit (Bruker). Emitted fluorescence was captured by a QUAD GaAsP PMT array (Bruker), managed by a QUAD GaAsP Controller (Bruker).

Environmental stability during live imaging was maintained using a custom aluminum heated stage and a specialized plate cover integrated with a CO_2_ tank and humidifier to preserve physiological temperature, humidity, and CO_2_ levels. Images were acquired through a Nikon 16X LWD water-immersion objective (0.80 NA, 3.0 mm WD) at a resolution of 1024 X 1024 pixels. The fields of view (FOV) ranged from 829 X 829 µm (1X zoom) to 51.8 X 51.8 µm (16X zoom), with frame rates standardized between 15.023 and 15.11 Hz.

For functional GCaMP imaging, two to five z-planes were acquired per organoid to sample distinct neuronal populations. Each recording consisted of 4,507 consecutive frames, with 4–8 organoids imaged per cell line at each time point. To validate the contribution of GABAergic inhibitory signaling to the observed network synchrony, the GABA_A_ receptor antagonist Gabazine (SR-95531) (Tocris, 1262) was acutely applied to the imaging medium at a final concentration of 10 µM. Post-drug recordings were initiated 5–10 minutes following application to ensure stable steady-state blockade. For structural analysis, dendritic spine imaging was performed on 4–8 organoids per line, with two to five dendritic segments captured per organoid at a higher magnification (16X zoom), totaling 600–900 frames per segment to ensure high-resolution volumetric reconstruction.

### GCaMP Functional Activity Analysis in Organoid

The images acquired via PrairieView software (Bruker) were saved as TIF files to be pre- processed via Suite2p^59^. The pipeline began with rigid registration utilizing regularized phase correlations to correct for motion artifacts. Neuronal regions of interest (ROIs) were subsequently identified through a clustering algorithm that groups correlated pixels, supported by a low-dimensional decomposition to optimize computational efficiency. While the initial number of ROIs was determined automatically via a pixel-correlation threshold, we performed a rigorous manual validation of all assigned ROIs^59^. This secondary check ensured that each ROI was classified based on accurate anatomical location, characteristic neuronal morphology, and the presence of biologically plausible ΔF/F transients. ΔF/F is defined as the fluorescence change calculated after performing local neuropil subtraction to eliminate background contamination. The corrected fluorescence (F_corrected) was estimated to avoid signal contamination from the surrounding dense neuropil according to the following formula:

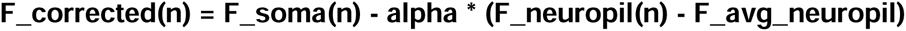

- **F_soma(n)**: Fluorescence of the soma at frame n.
- **F_neuropil(n)**: Fluorescence in the region less than 30 micrometers from the ROI border at frame n.
- **F_avg_neuropil**: The average neuropil fluorescence across all frames.
- **alpha**: A correction factor chosen to minimize the Pearson’s correlation coefficient between F_soma and F_neuropil.

The relative change in fluorescence for each neuron was then calculated as:

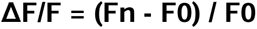

- **Fn**: The corrected fluorescence (F_corrected) for frame n.
- **F0**: The baseline fluorescence, defined as the first mode of the corrected fluorescence density distribution across the entire time series.

To quantitatively evaluate functional connectivity and network dynamics, we developed a custom Python pipeline to analyze ΔF/F and estimated spike data extracted from Suite2p (<u>Organoid_GCaMP_pipeline</u>). To ensure the inclusion of only high-quality biological signals, a two-stage filtering process was implemented: (1) a basic signal quality filter was applied to remove ROIs with low peak amplitudes (bottom 10th percentile) or atypical variance, and (2) an event-based signal-to-noise ratio (SNR) filter was utilized to exclude cells lacking distinct calcium transients, using an adaptive threshold based on the median absolute deviation (MAD) of the signal. For the surviving ROIs, functional connectivity was determined by calculating the maximum cross-correlation with time lags to account for slight temporal offsets in neuronal firing.

To evaluate the cross-correlation of neuronal activity among surviving ROIs and correct for chance-level correlations, a circular shuffling procedure was implemented across 1,000 iterations. For each ROI, the spike trace was independently shifted by a random circular frame offset to generate a null distribution of cross-correlation values. For each recording, the mean cross-correlation value across all ROI pairs was calculated, and the corresponding mean chance correlation derived from the shuffling baseline was subtracted from it to yield the corrected cross-correlation metric. For matched pre- and post-treatment recordings, the binary ROI filtering mask established during the baseline pre-treatment condition was locked and directly applied to all corresponding post-treatment recordings, ensuring that identical neuronal ensembles were tracked longitudinally.

### Dendritic Spine Morphology Analysis in Organoid

Two-photon image stacks were acquired via PrairieView software (Bruker) and converted into multi-page TIF files. The raw TIF files were first preprocessed via a specialized MATLAB-based framework (<u>A ProcessTimeSeries.m</u>). This step involved extracting the fluorescent time series and performing X-Y image registration to correct for any lateral movement during the recording sessions. Following motion correction, an activity map was generated for each recording to facilitate the identification of regions of interest (ROIs). This was achieved by calculating either the local correlation or a modified ΔF/F based on kurtosis to highlight pixels with significant fluorescence signals over time^60–62^.

Dendritic segments were processed using a different pipeline (<u>Organoid_Dendritic_Spine_ID.m</u>) configured with parameters specifically optimized for organoid-derived neuronal architectures. To prepare the data for analysis, each of the raw TIF image were first converted into average projection images and saved as PNG files using ImageJ. Within the custom MATLAB pipeline, these images underwent binarization via manual thresholding to ensure accurate signal representation. The classification process involved segmenting each dendritic spine to confirm the presence or absence of a spine neck^60^. Utilizing the binarized images, each identified spine was then categorized according to the organoid-specific spine classification criteria integrated into the pipeline (see Supplementary Table 2 for representative morphological thresholds)^60–62^. Given the inherent challenges of imaging organoid tissues, such as low signal-to-noise ratios and background interference, we employed a hybrid analysis approach. While initial spine identification was performed via automated scripts, all data underwent extensive manual inspection to correct for potential artifacts or weak signal intensity. This rigorous curation ensured that only clearly identifiable spines were included in the final density and maturation counts.

## RESULTS

### Developmental Anatomy of WS and Dup7 Organoids

Neural rosettes are 3D clusters of neural stem cells that resemble the radial arrangement of the developing embryonic neural tube in which precursor cells proliferate, differentiate, and migrate outward to form cortical layers that include specialized neurons, oligodendrocytes, and astrocytes^25^. In this role, their number and structure provide insight into the timing and nature of early cortical development. The developmental growth of organoids including the presence and subsequent maturational loss of neuroepithelial rosettes mark key stages of cortical differentiation^25,63^. To systematically investigate these neurodevelopmental dynamics, we established an experimental pipeline spanning early structural and histological characterization to late-stage dendritic spine profiling and longitudinal two-photon calcium imaging of functional network (Fig. 1A). We analyzed organoid cross-sectional area, and their neurodevelopmental patterning as measured by rosette number and their thickness as a function of genotype and age. To assess early tissue expansion across genotypes, we quantified area in four samples per line from embryoid bodies (EBs) to the onset of organoid formation every five days from day 1 to day 55 (Supplementary Fig. 1). The largest diameter was measured as they grew larger and more spherical from day 30 to day 150 and converted to cross-sectional area for comparison (Supplementary Fig. 2).

**Figure 1.**
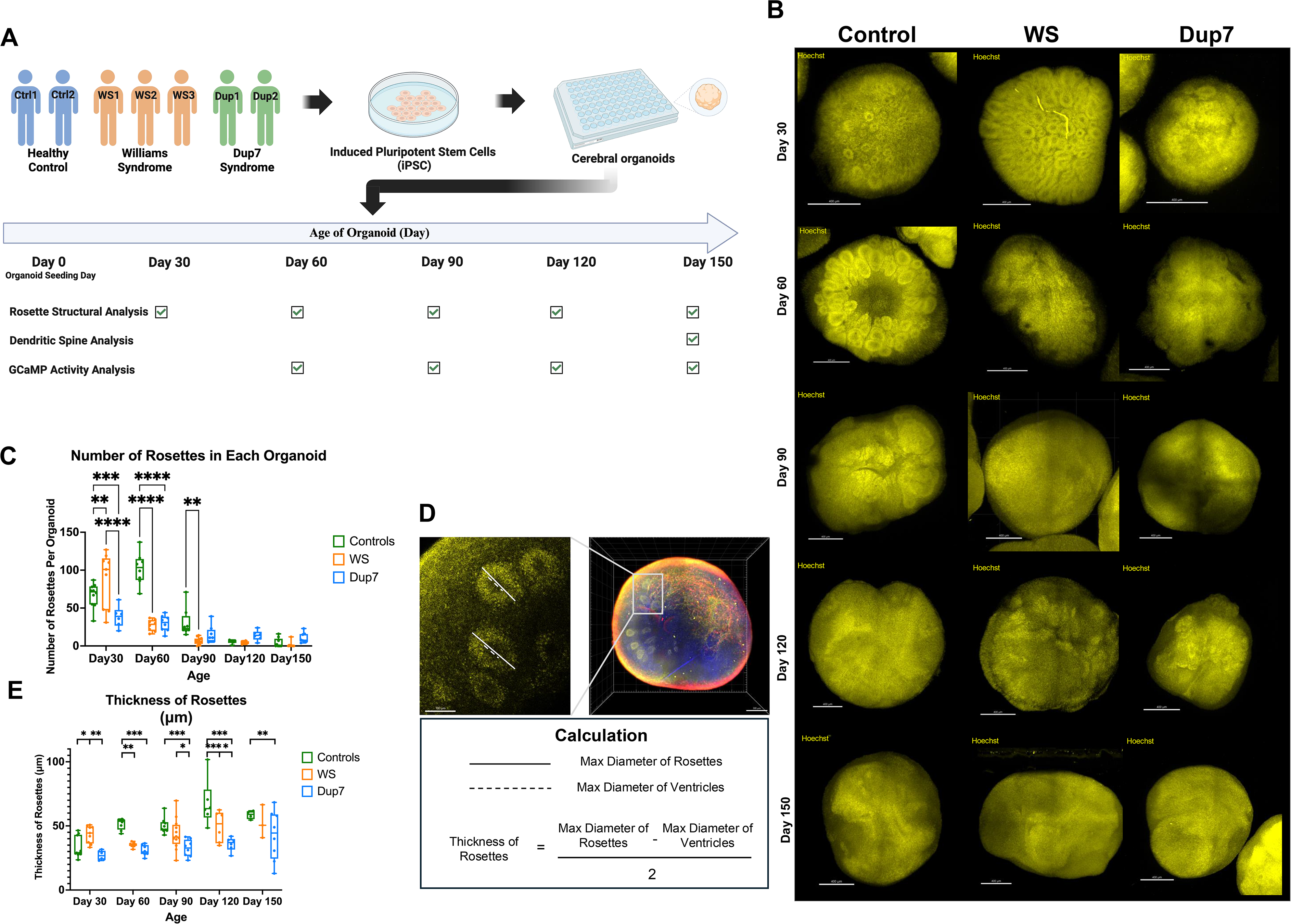
Longitudinal characterization of rosette formation and morphometric analysis. (**A)** Experimental overview: A total of 7 reprogrammed iPSC lines, derived from 7 individuals, were differentiated into cerebral organoids and harvested at longitudinal time points for multifaceted analyses, including GCaMP6m calcium imaging, rosette structural profiling, and dendritic spine quantification. Created in BioRender. Hwang, I. (2026) https://BioRender.com (**B**) Representative images of cortical rosettes in cleared organoids across all genotypes (Control, WS, and Dup7). Tissues were stained with Hoechst and imaged at Day 30, 60, 90, 120, and 150 to capture the developmental trajectory of progenitor patterning. (**C**) Schematic of rosette identification and morphometric quantification. Middle: Representative image of a cleared organoid. Left: Magnified view illustrating the criteria for identifying individual rosettes and measuring their thickness. Right: Geometric equation used for the calculation of rosette thickness. **(D)-(G)** N = 36–43 organoids per genotype integrated from 2–3 cell lines across 2–4 independent differentiation batches. (**D**) Longitudinal tracking of rosette number. Statistical assessment of the number of rosettes formed per organoid for each genotype. Each data point represents the total rosette count in an individual organoid. Longitudinal quantification of total rosette numbers derived from three 7q11.23 genotypes evaluated across five sequential ages (N = 120 total values). Ordinary Two-way ANOVA revealed highly significant main effects for Age (F(4, 105) = 80.93, P < 0.0001) and Genotype (F(2, 105) = 21.05, P < 0.0001). A highly significant Interaction effect (F(8, 105) = 18.84,P < 0.0001; accounting for 22.80% of total variation) statistically confirms that the temporal trajectories of rosette morphogenesis diverge significantly based on gene dosage. Data were integrated from 6–12 organoids per genotype, derived from 2–3 independent cell lines across 2–4 distinct differentiation batches. Boxes indicate the 25th–75th percentiles; whiskers represent the minimum to maximum values. **(E)** Spatiotemporal analysis of rosette thickness. Statistical quantification of individual rosette thickness across the 150-day period. Each data point represents the average thickness of rosettes per organoid, serving as the single representative value per organoid. Ordinary two-way ANOVA revealed significant main effects for both Genotype (F(2,95) = 5.34, P = 0.006) and Time (F(4,95) = 15.46, P < 0.001), along with a significant Genotype × Time interaction (F(8,95) = 3.83, P = 0.001; overall model R² = 0.58). Pairwise genotype comparisons were performed within each timepoint using estimated marginal means. Data represent mean ± SEM; *P < 0.05, **P < 0.01, ***P < 0.001.

Control EBs displayed a linear growth trajectory from day 1 through day 55, reflecting stable early neuroectodermal expansion (Supplementary Fig. 1). In contrast, WS EBs exhibited accelerated growth around day 20, at which time these organoids deviated markedly from the normative growth pattern (Supplementary Fig. 1). Dup7 EBs showed the slowest growth rate and the smallest EB sizes from day 1 through day 55. By day 90 and through day 150, all groups plateaued at statistically identical sizes (Supplementary Fig. 2). Representative confocal images of whole-mount organoids revealed clear temporal and genotype-specific differences in neural rosette morphology acros development (Fig. 1B and Supplementary Fig. 3). Under control conditions, as organoids mature, the number of rosettes decreases^63^. In control organoids, the mean number of rosettes per organoid peaked up to 102.0 at day 60 before undergoing a sharp reduction to 30.38 by day 90 and eventually plateauing at 4.11 by day 150. WS organoids exhibited an accelerated trajectory, reaching their mean rosette number (87.56) much earlier at day 30, followed by a decline to 27.25 at day 60 and 1.75 by day 150. Dup7 organoids displayed the most suppressed expansion pool size, showing a mean of 38.50 rosettes at day 30, which gradually decreased to 29.75 at day 60 and maintained a higher baseline of 10.63 at Day 150 (Fig. 1C). This result suggested that tracking rosettes over development may be informative with regard to genotype. Intriguingly, while the overall organoid size at day 30 showed no significant difference across genotypes (Supplementary Fig. 2), our statistical analysis revealed highly divergent profiles in rosette number at this time point. This divergence indicates that the internal structural remodeling occurring within the organoid proceeds independently from the overall physical growth and volumetric expansion of the organoid.

To evaluate these structural features, rosette number and rosette thickness were analyzed independently as a function of genotype (Control, WS, Dup) and age (Day 30, 60, 90, 120, 150), using two-way ANOVAs. We measured rosette thickness as the diameter of individual rosettes minus the diameter of their central lumens divided by two (Fig. 1D). Thickness measurements were averaged within each organoid to yield a single representative value as the unit of analysis. For total rosette number (Fig. 1C), the two-way ANOVA revealed highly significant main effects for age (F(4, 105) = 80.93, P < 0.0001) and genotype (F(2, 105) = 21.05, P < 0.0001), as well as a highly significant genotype x age interaction effect (F(8, 105) = 18.84, P < 0.0001). For rosette thickness (Fig. 1E), a two-way ANOVA similarly revealed significant main effects of both genotype (F(2,95) = 5.34, P = 0.006) and age (F(4,95) = 15.46, P < 0.001), along with a significant genotype × age interaction (F(8,95) = 3.83, P = 0.001). Rosette number and thickness exhibited highly parallel phenotypic trends across development. At day 30, WS organoids displayed a pronounced, accelerated peak in total rosette number (Fig. 1C) that tightly synchronized with a significantly greater rosette thickness compared to both control and Dup7 cohorts (Fig. 1E). Control organoids reached their peaks later, at the day 60 time point, displaying a concurrent maximum in both rosette number and thickness. Following these peaks, both metrics showed a progressive decline. Control rosettes maintained a higher rosette number and remained significantly thicker than Dup7 across subsequent stages. By day 150, the structural differences attenuated: WS and Dup7 groups no longer differed significantly in either number or thickness, and control and WS rosette thicknesses were not significantly different, while Dup7 rosettes remained significantly thinner than those of controls (Fig. 1E). Data were integrated from 6–12 organoids per genotype, derived from 2–3 independent cell lines across 2–4 distinct differentiation batches (Fig. 1C and 1E). Together, these analyses demonstrate that rosette number and thickness follow parallel development profiles, statistically confirming that the temporal trajectories diverge significantly by genotypes.

To ensure that these size variations did not stem from localized necrosis or abnormal developmental arrest, we performed routine quality checks by harvesting organoids every 30 days starting from day 30. Histological and marker evaluations confirmed that organoids across all genotypes were growing and maturing within normative physiological ranges throughout the culture period (Supplementary Fig. 4)

In summary, these results reveal that 7q11.23 CNVs reshape early neural progenitor dynamics and spatial organization. The observed perturbations in the timing and capacity of progenitor expansion underscore a profound shift in the early developmental blueprint. These early-stage disruptions establish a compromised foundation that likely drives the subsequent failures in functional circuit integration and the aberrant synaptic maturation seen at later time points. By disrupting the timing and inherent capacity for cortical maturation, 7q11.23 dosage imbalances dictate the divergent neurodevelopmental trajectories that characterize the WS and Dup7 phenotypes.

### Dup7 organoids exhibit higher dendritic spine density and more mature dendritic spine morphologies than WS

The dysmorphic rosette organization and altered proliferation timelines observed suggested potential long-term consequences on neuronal and synaptic maturation. We performed high-resolution two-photon imaging of dendritic spine morphology at Day 150 (Fig. 1A and Fig. 2A). Given that dendritic spine morphology has not been extensively quantified or categorized in cerebral organoid models^28,29^, we first established a specialized identification pipeline and classification criteria tailored for organoid-derived neuronal architectures (Supplementary Table 2). Our quantitative analysis utilizing this framework revealed that Dup7 organoids exhibited a significantly higher dendritic spine density compared to WS, while no significant differences were observed comparing the control to WS or Dup7 (Fig 2B). This finding suggests that the Dup7 genotype has either a more precocious establishment of synaptic programs or failure to prune synapses compared to the WS group. Overall spine densities remained relatively low (< 1 spine per 10 µm; Fig. 2B). Organoids across all three genotypes manifested a diverse repertoire of spine morphologies—including filopodia, thin, stubby, and mushroom types, recapitulating the structural diversity seen *in vivo* (Supplementary Fig. 5). The identification of these distinct maturational stages provides definitive evidence of functional synaptogenesis within the organoid model. Furthermore, these results underscore that cerebral organoids can serve as a platform for investigating dendritic spine dynamics and synaptic development.

**Figure 2.**
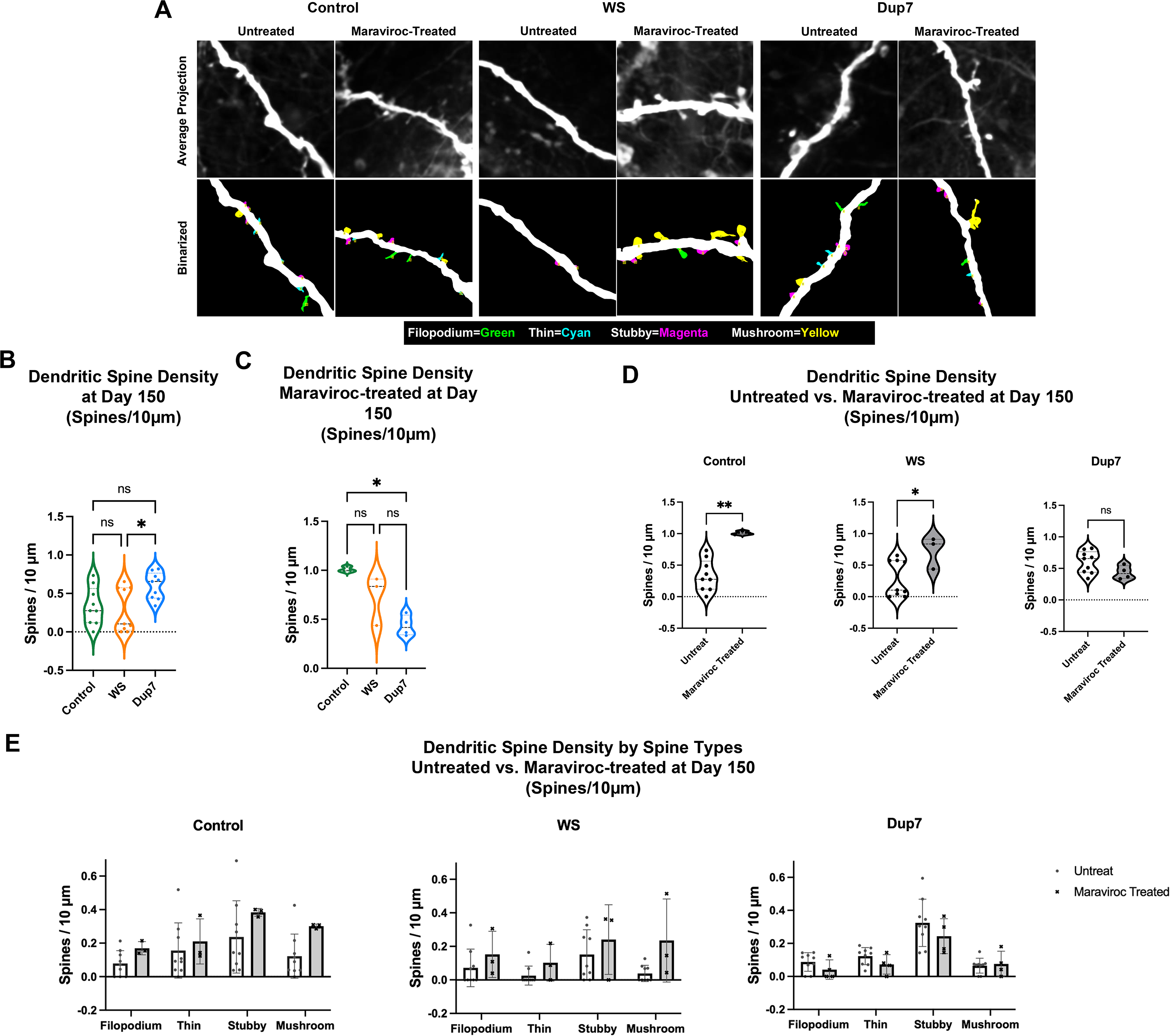
Morphological characterization of dendritic spines and therapeutic rescue via Maraviroc. (**A**) Representative raw images of dendritic spines. Comparison of Control, WS, and Dup7 organoids at Day 150 in untreated and Maraviroc-treated conditions. Top row: Average projection of raw two-photon images. Bottom row: Corresponding binarized masks with identified spine types color-coded (Filopodia: green; Thin: cyan; Stubby: magenta; Mushroom: yellow). (**B**) Quantitative analysis of dendritic spine density at Day 150. Comparison of baseline spine density (spines per 10 μm) across genotypes. Statistical significance was determined by an ordinary one-way ANOVA non-parametric test (p = 0.0335). Each data point represents the average spine density per organoid. (**C**) Impact of Maraviroc treatment on total dendritic spine density. Comparative analysis of spine density at Day 150 following Maraviroc treatment. Statistical significance was determined by an ordinary one-way ANOVA non-parametric test (p = 0.01). Each data point represents the average spine density per organoid. (**D**) Comparative efficacy of Maraviroc on spine density. Statistical comparison between untreated and Maraviroc-treated groups within each genotype using a non-parametric one-tailed t-test. Control (p = 0.0045), WS (p = 0.0500), and Dup7 (p = 0.0741, ns). (**E**) Subtype-specific responses to Maraviroc. Statistical assessment of density changes across individual spine subtypes following Maraviroc treatment, analyzed via non-parametric multiple t-tests across all genotypes and spine subtypes at Day 150.

### Maraviroc promotes structural synaptic plasticity in genotype-dependent manner

Maraviroc is a potent antagonist of C-C chemokine receptor 5 (CCR5), a seven-transmembrane G protein-coupled receptor traditionally recognized for its role in immune signaling and as a therapeutic target for HIV/AIDS^34–36,64^. Recent studies have identified CCR5 as a key negative regulator of synaptic plasticity, whose inhibition has been shown to enhance MAPK and CREB signaling, thereby promoting cortical and hippocampal-dependent learning and memory^35,36,64^. To establish a biological rationale for this pathway, we confirmed robust and uniform CCR5 expression across all three genotypes at Day 150 (Supplementary Fig. 6). Given that CCR5 antagonism via Maraviroc has been reported to facilitate neural repair and dendritic spine turnover in other models^35,36^, we investigated whether these synaptoplastic effects could be recapitulated in human iPSC-derived organoids.

Beginning at Day 120, organoids were treated with 200 ng/mL of Maraviroc every other day for 30 days^58^. To visualize these structural changes, projection images were generated by averaging all frames from the raw two-photon stacks. These images were then binarized using appropriate intensity thresholds to allow for the precise categorization of spine morphologies, with each distinct spine type assigned a specific color for identification. As shown in the representative dendrite images (Fig. 2A), quantification of these morphological features revealed that Maraviroc treatment significantly increased total dendritic spine density in both control and Williams Syndrome (WS) organoids (Fig. 2C and 2D). Beyond the total count, an increasing trend was observed across all individual spine subtypes within these two genotypes, encompassing the full morphological repertoire from filopodia to mature mushroom spines (Fig. 2E). Notably, this pro-synaptogenic effect was absent in Dup7 organoids, further highlighting a genotype-specific response to CCR5 antagonism. Considering the well-characterized neurocognitive phenotypes associated with WS^1^, these findings suggest that Maraviroc- mediated CCR5 inhibition may represent a possible therapeutic strategy for modulating synaptic deficits in this population.

### WS syndrome cerebral organoids exhibit delayed and less persistent synchronized bursting, while synchrony is abolished in Dup7

We next sought to capture the functional readout of the 7q11.23 CNVs at the ensemble level. To examine the developmental emergence of coordinated neuronal activity, we performed two-photon calcium imaging of control, WS, and Dup7 cerebral organoids from day 60 to day 150 (Fig. 1A). Spontaneous calcium transients were observed in all organoids across all genotypes and time points. However, the degree and timing of network firing synchrony varied significantly by genotype (Fig. 3A). In control organoids, synchronous network firing was evident as early as day 60 and persisted through day 150 (Fig. 3A, Video S1). In contrast, such synchrony was not observed in WS until day 90 and diminished again around day 120. Dup7 organoids showed the most profound impairment, exhibiting minimal to no synchronous activity across all time points examined (Fig. 3A, Video S2-S3).

**Figure 3.**
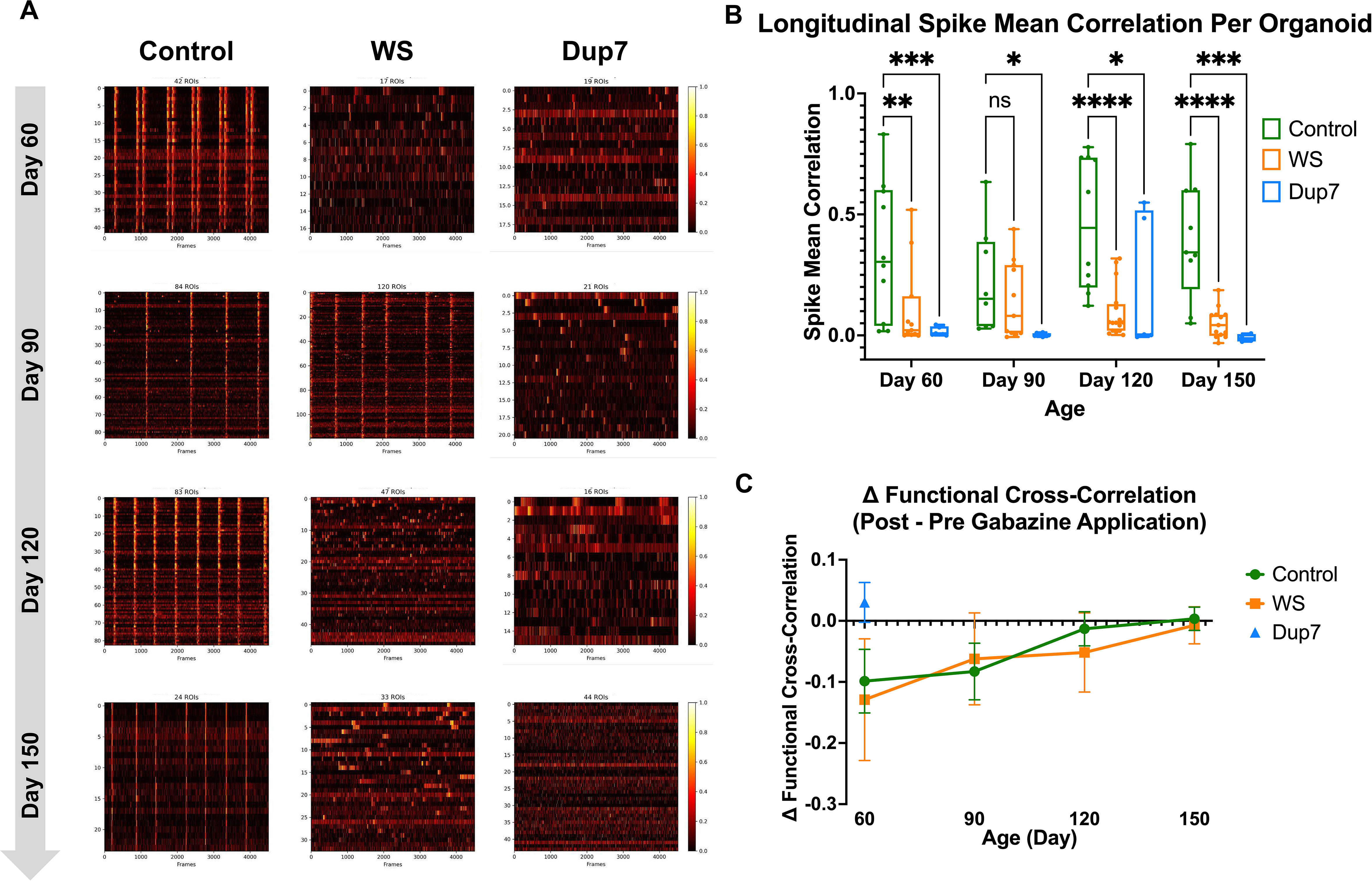
Functional network alterations in 7q11.23 CNV organoids. **(A) Corresponding functional cross-correlation matrices for Control, WS, and Dup7 organoids across the 150-day developmental trajectory.** **(B) Statistical quantification of spontaneous spike mean correlation.** Each data point represents an individual organoid, with mean spike cross-correlation values calculated across two-photon imaging sessions (4,507 frames per session) to account for nested variance. Data were integrated from 4–16 organoids per genotype, sourced from 2–3 independent cell lines across 2–4 distinct differentiation batches. Data are presented as box-and-whisker plots, where the middle horizontal line represents the median, the box bounds indicate the 25th and 75th percentiles, and the whiskers encompass the full range from minimum to maximum values (min-to-max). Statistical evaluation via ordinary two-way ANOVA revealed a highly significant main effect of genotype (F(2, 113) = 21.51, P < 0.0001), accounting for 24.81% of the total variation. In contrast, no significant main effect was observed for age (F(3, 113) = 0.6188, P = 0.6042) or the genotype x age interaction (F(6, 113) = 2.746, P = 0.0158, accounting for 9.504% of total variation; total n = 125). Adjusted P values are indicated as follows: * P < 0.05, P < 0.01, *** P < 0.001, **** P < 0.0001, and ns, not significant. **(C) Line graph displaying the mean change in spike cross-correlation following Gabazine application.** Calculated as post-Gabazine cross-correlation minus pre- Gabazine cross-correlation) across developmental time points. Negative values indicate a reduction in functional network synchronization in response to GABA_A_ receptor blockade. Data are presented as mean standard error of the mean (SEM) for each genotype at each time point.

Functional cross-correlation matrices generated from each recording closely paralleled these observations (Supplementary Fig. 7). Control organoids displayed strong correlations in neuronal firing patterns indicative of robust network coupling. WS organoids showed increased cross-correlation compared to their own earlier time point at day 60 and corresponding to their brief period of hyper-synchrony—while Dup7 organoids remained weakly correlated at all stages (Supplementary Fig. 7).

We calculated the proportion of organoids exhibiting robust synchronous firing events for each genotype at every time point (Supplementary Table 3). Control organoids consistently exhibited a higher proportion of synchronously firing organoids compared to both WS and Dup7 across all analyzed developmental stages (Supplementary Table 3), except at day 90, the proportion of synchronous organoids in WS exceeded control (Supplementary Table 3). In terms of functional correlation strength, control organoids exhibited significantly higher spike mean correlation than both WS and Dup7 at days 60, 120, and 150, reflecting a sustained and strengthening network throughout development (Fig. 3B). At day 90, the specific developmental window where WS organoids manifested transient synchrony—no significant difference in correlation was observed between WS and controls (Fig. 3B). Dup7 organoids displayed a near-absence of synchronous activity across all stages (Fig. 3B). No significant differences between WS and Dup7 were detected at other time points (Fig. 3B).

Of note is that in the developing human cortex, early spontaneous synchronous and oscillatory network activity—often referred to as spontaneous synchronized bursts or early network oscillations—is essential for guiding synaptic refinement, establishing excitatory– inhibitory balance, and shaping the emergent architecture of functional neuronal circuits^38,40,47,65–67^. The early onset and prolonged presence of synchrony in control organoids are therefore consistent with normal maturation of cortical networks. In contrast, the delayed and transient synchrony observed in WS organoids suggests a disruption in circuit assembly. The near-complete absence of synchronous activity in Dup7 organoids across all imaged time points indicates a profound impairment in functional network integration. These results demonstrate that 7q11.23 dosage alteration significantly shifts the trajectory of early network maturation, with potential downstream consequences for synaptic development and the emergence of higher-order circuit functions. Ultimately, these functional deficits in network coordination may contribute to the diverse cognitive and neurological symptoms^1,2,7^ observed in clinical populations with 7q11.23 copy number variations.

### Differential Effects of Gabazine on Synchronous Network Firing in 7q11.23 syndromes

In the immature neural network, blocking GABA_A_ receptors with the highly potent, selective antagonist gabazine of GABA_A_ abolishes or scatters the physiological, spontaneous synchronous network bursting because GABA acts as a primary depolarizing driver during early cortical development^39,68–70^. To evaluate whether the synchronized activities in our organoids can be interrupted by a GABA_A_ antagonist we quantified the cross-correlation coefficients before and after gabazine application from day 60 to day 150 (Fig. 3C and Supplementary Fig. 8). Consistent with the literature (Ben Ari et al. 1989; Garaschuk et al. 1998; Khalilov et al. 1999; Sipila et al. 2005), control organoids exhibited a substantial reduction in cross-correlation coefficients following gabazine application (Fig. 3C). In controls, the spontaneous cross-correlation remained high throughout all developmental time points(Fig. 3C). Representative changes in network activity following gabazine treatment are shown in Video S4–S5. At day 60, the reduction in cross-correlation following gabazine treatment was most pronounced in WS organoids, whereas control organoids exhibited the greatest reduction at day 90 (Fig. 3C). Analysis for Dup7 organoids was restricted to day 60, as this was the only developmental stage at which Dup7 displayed baseline spontaneous synchrony (Fig. 3B-C). As occurs in all organoid experiments, occasional paradoxical responses are observed; indeed, one Dup7 organoid showed a more well-defined synchronous activity after gabazine treatment that remains unexplained (Fig. 3C). Overall, these findings demonstrate differential temporal sensitivity to GABA_A_ receptor blockade among control, WS, and Dup7 organoids during network maturation.

## Discussion

Syndromes with hemizygous deletions or duplications of 25-27 genes in the 7q11.23 region lack a clear explanation of how the genotype is related to the complex phenotype. Inferring the collective impact of the entire 7q11.23 locus from single gene deletion experiments has severe limitations. To date, research on the 7q11.23 region has largely centered on gene-specific investigations, focusing on individual candidates such as BAZ1B, GTF2I, FZD9, STX1A, TTR and LIMK1^2,4,5,13,19,20,22–24,27,71,72^. These studies have demonstrated that specific genes can contribute to the neurodevelopmental phenotype often centered around altered time course patterns of gene expression suggesting either developmental delay or acceleration^19^. Extrapolating additional specific phenotypic features could potentially link these features to specific gene dosage effects. From brain organoids carrying 7q11.23 deletion or duplication, numerous new phenotypic features can be inferred including organoid rosettes, spines and circuitry. The observation that Maraviroc (a CCR5 antagonist) significantly increased dendritic spine density in WS organoids provides a promising therapeutic lead. Given that CCR5 signaling is a known modulator of neuronal plasticity^31,34–36,64^, our results suggest that pharmacological intervention can recalibrate the structural imbalances inherent in WS.

Asynchronous activity has been increasingly recognized across various neurodevelopmental models, particularly in the context of autism^73,74^. Recent work demonstrated that haploinsufficiency of three autism-related genes (SUV420H1, ARID1B, CHD8) resulted in asynchronous development of GABAergic neurons and deep-layer excitatory projection neurons acting through distinct molecular pathways^73^. Based on these molecular developmental differences, the authors looked at bursting activity, a typical neurophysiological feature of brain organoids present within a developmental window^37^. Presumably the asynchronous cellular development contributed to altered burst duration and, they concluded that in controls, a lower sensitivity to the application of NBQX, an antagonist of non-NMDA glutamate receptors, suggested autism mutant cells were more mature^73,74^. Because early synchronous network activity is crucial for early neuronal survival and integration^75–77^, the shortened or absent synchronous firing periods in WS and Dup7 organoids may further exacerbate this reduction in brain volume and compromise the integrity neuronal maturation.

These studies suggested that features of bursts would be informative among cases of 7q11.23 CNVs especially when performed over a longitudinal developmental period of 150 days. From this extended observation window, we observed the full trajectory of maturation. Our rosette analysis reveals a reduction in the duration of the proliferative phase alongside compromised integrity of proliferative capacity in both WS and Dup7 organoids, which we speculate may underlie the reduced brain volume observed in WS and Dup7 patients^1^. Furthermore, the abnormal patterns of rosette formation may result in the underlying asynchronous development of neuronal populations^73,74^ However, the autistic features of Dup7 cases is associated with a more severe phenotype than the ASD haploinsufficiency genes^73^ or PTEN mutant genes^74^. In the analysis of our data inferences that conclude accelerated or delayed maturation seem to be a simplification. Synchronous firing indicative of network activity is altered in both WS and Dup7 and these alterations have qualitative features more complex than can be explained by a time-shifted developmental clock. GABAergic neurons likely contribute based on the effects gabazine. The relatively late emergence of inhibitory neurons, albeit in small numbers in this organoid system^19^ in both WS and Dup 7 may contribute to the longitudinal changes in synchrony.

**Table 1.**
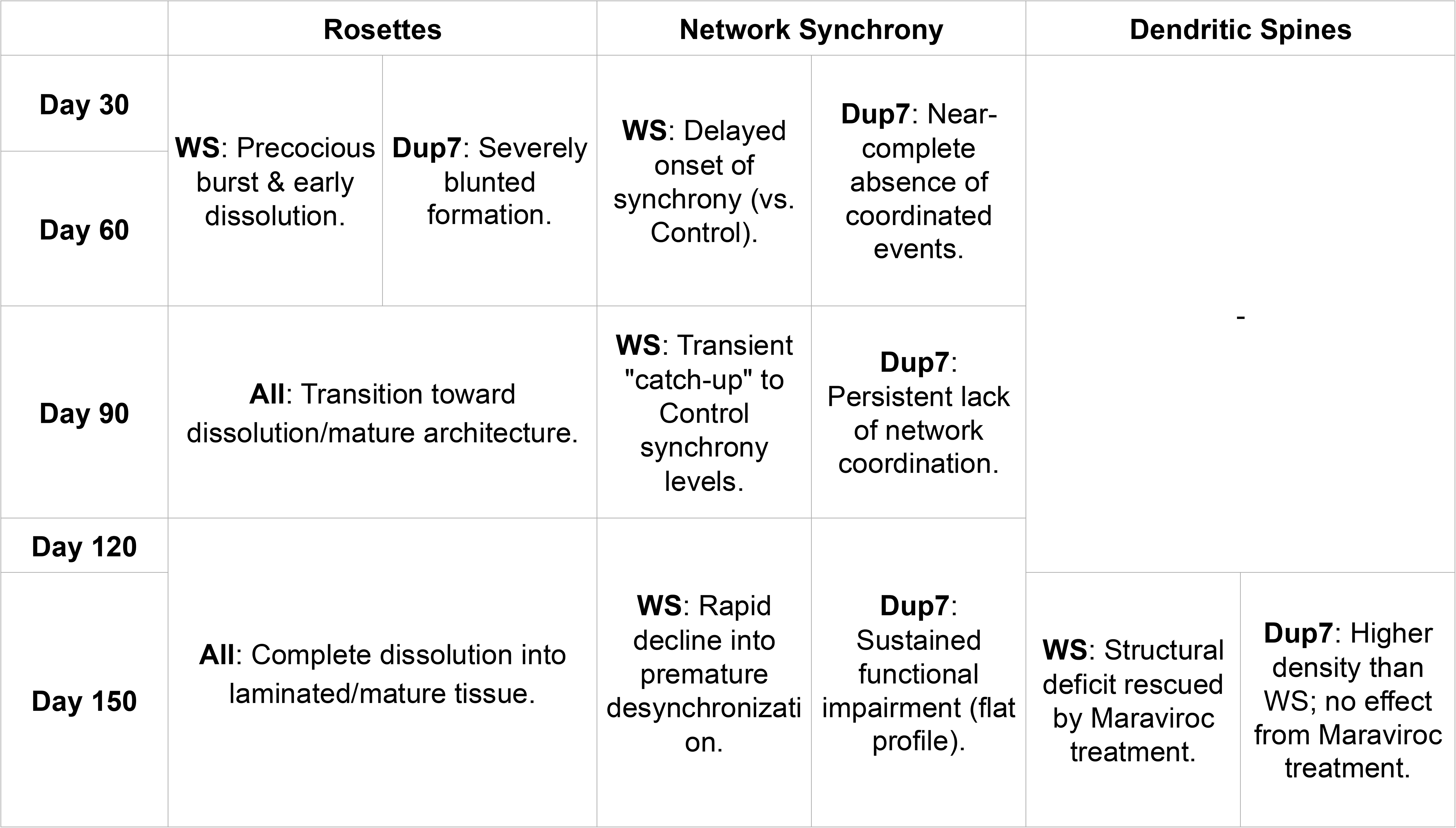
Integrated Summary of 7q11.23-Dependent Developmental Milestones. A longitudinal comparison of progenitor organization, network functional maturation, and structural synaptogenesis across Control, WS, and Dup7 genotypes. Rows represent key developmental time points, while columns categorize findings by biological modality: neuroepithelial rosette dynamics, GCaMP-based network synchrony, and dendritic spine morphology. Each cell highlights the primary phenotypic deviation or hallmark specific to each genotype. Statistical significance and detailed quantification for each modality are provided in Figure 1, Figure 2 and Figure 3, and associated Supplementary Materials.

## Supporting information

CNV info

## Acknowledgments

We deeply thank the patients and their families for making this study possible. We extend our gratitude to our collaborators for their valuable insights and contributions. We thank Tadeo Thompson for reprogramming the WS3 line and Caitlin Loo for the initial pluripotency characterization at The Hospital for Sick Children. Schematics and the graphical abstract were created with BioRender.com. This work was supported by the National Institutes of Health (NIH 2R01 AG056058 to K.S.K.) and John Douglas French Alzheimer’s Foundation.

## Author Contributions

The experiment was designed by I.H. and K.S.K. Methodologies were established by I.H., J.S.Y., M.J.G., and I.T.S. All the experiments and data collection were performed by I.H., with experimental assistance from C.C., M.C.A., D.C.C., J.A.U., A.H., and A.N. Key resources and sample lines were provided by A.A., M.B., J.E., and L.R.O. All the data were analyzed and visualized by I.H. and J.S.Y. The project was supervised and funded by K.S.K. The manuscript was written by I.H. and K.S.K. with feedback from all the authors.

