## Supplementary Fig and Tables for "Disrupted Brain Organoid Circuitry, Structural Organization, and Spine Morphology in 7q11.23 Copy Number Variant Syndromes"

#### Supplementary Table 1

| Cell line | Original Code | Gender | Age | Position of Deletion or Duplication within 7q11.23 Locus |
|---|---|---|---|---|
| **Ctrl1** | F12442.4 | Female | 30 | - |
| **Ctrl2** | KOLF2.1J | Male | 55-59 | - |
| **WS1** | - | Female | 9 | Deletion: 73,303,743 - 74,200,000 |
| **WS2** | - | Female | 5 | Deletion: 72,700,000 – 74,200,000 |
| **WS3** | WS A | Male | 11 months | Deletion: 73,303,743 – 74,200,000 |
| **Dup1** | Dup A | Female | Unknown | Duplication: 73,303,743 – 74,200,000 |
| **Dup2** | - | Female | 11 | Duplication: 72,930,012 – 74,200,000 |

#### Supplementary Table 2

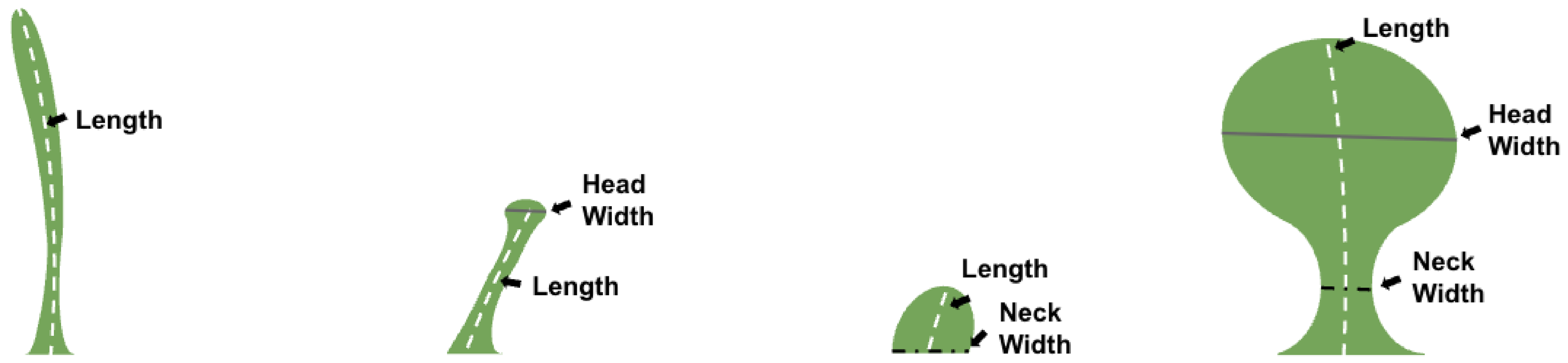


| Filopodium | Thin | Stubby | Mushroom |
|---|---|---|---|
| Length ≥ 3.5 um | 0.6 ≥ Head / Length ≥ 0.35 | 1.2 ≥ Neck / Length ≥ 0.6<br>6 um ≥ Length ≥ 0.5 um | Neck / Head ≤ 0.5 |

#### Supplementary Table 3

| Proportion of Synchronously Firing Organoids | | | | | | | | | |
|---|---|---|---|---|---|---|---|---|---|
| | Control | | | WS | | | Dup7 | | |
| | Total Active Organoids | Total Synchronous Organoids | Synchronous Proportion (%) | Total Active Organoids | Total Synchronous Organoids | Synchronous Proportion (%) | Total Active Organoids | Total Synchronous Organoids | Synchronous Proportion (%) |
| **Day 60** | 7 | 10 | 70 | 6 | 11 | 55 | 3 | 8 | 38 |
| **Day 90** | 5 | 8 | 63 | 7 | 11 | 64 | 1 | 9 | 11 |
| **Day 120** | 9 | 10 | 90 | 8 | 16 | 50 | 2 | 5 | 40 |
| **Day 150** | 7 | 9 | 78 | 3 | 13 | 23 | 1 | 4 | 25 |

**Supplementary Figure 1**

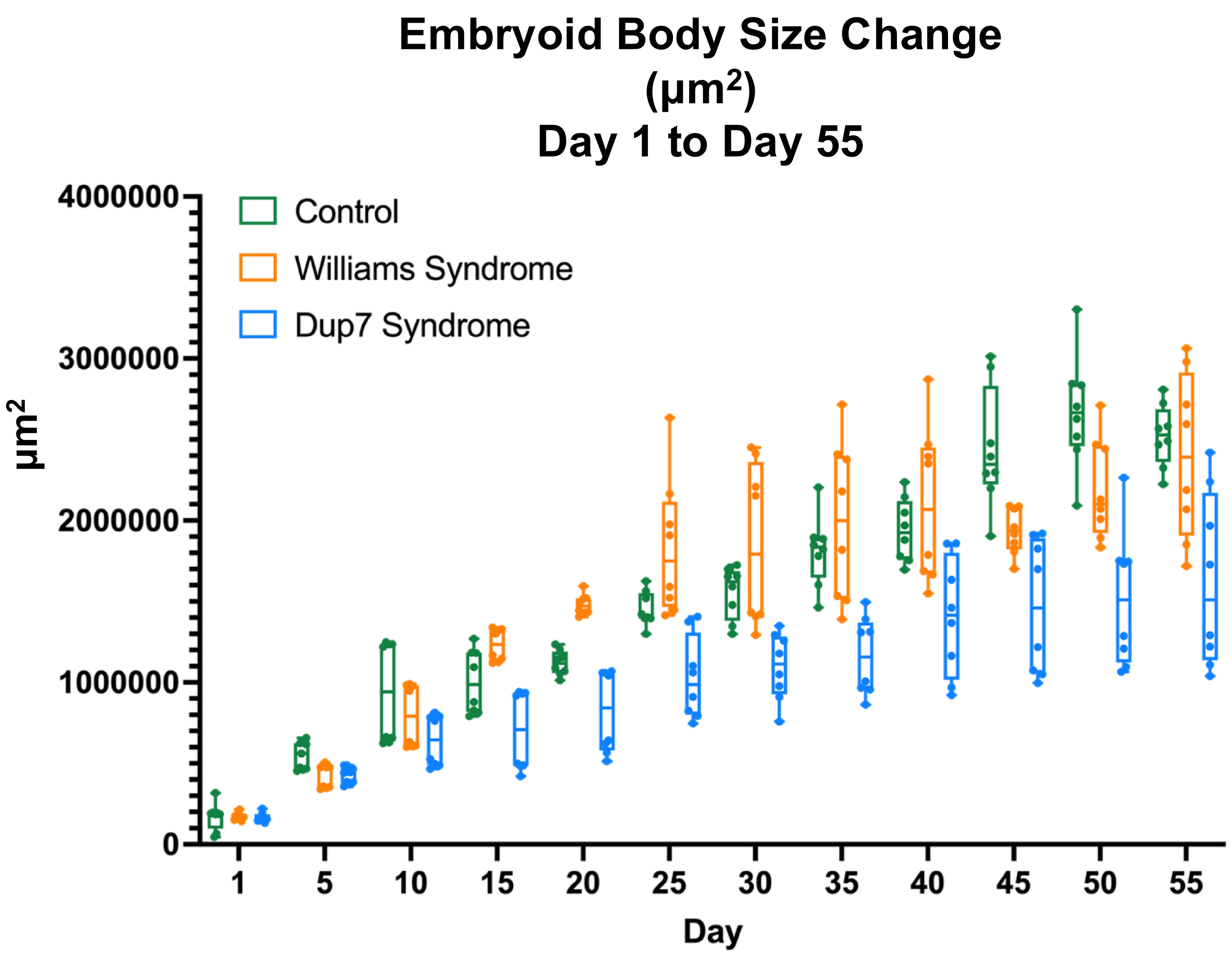


**Supplementary Figure 2**

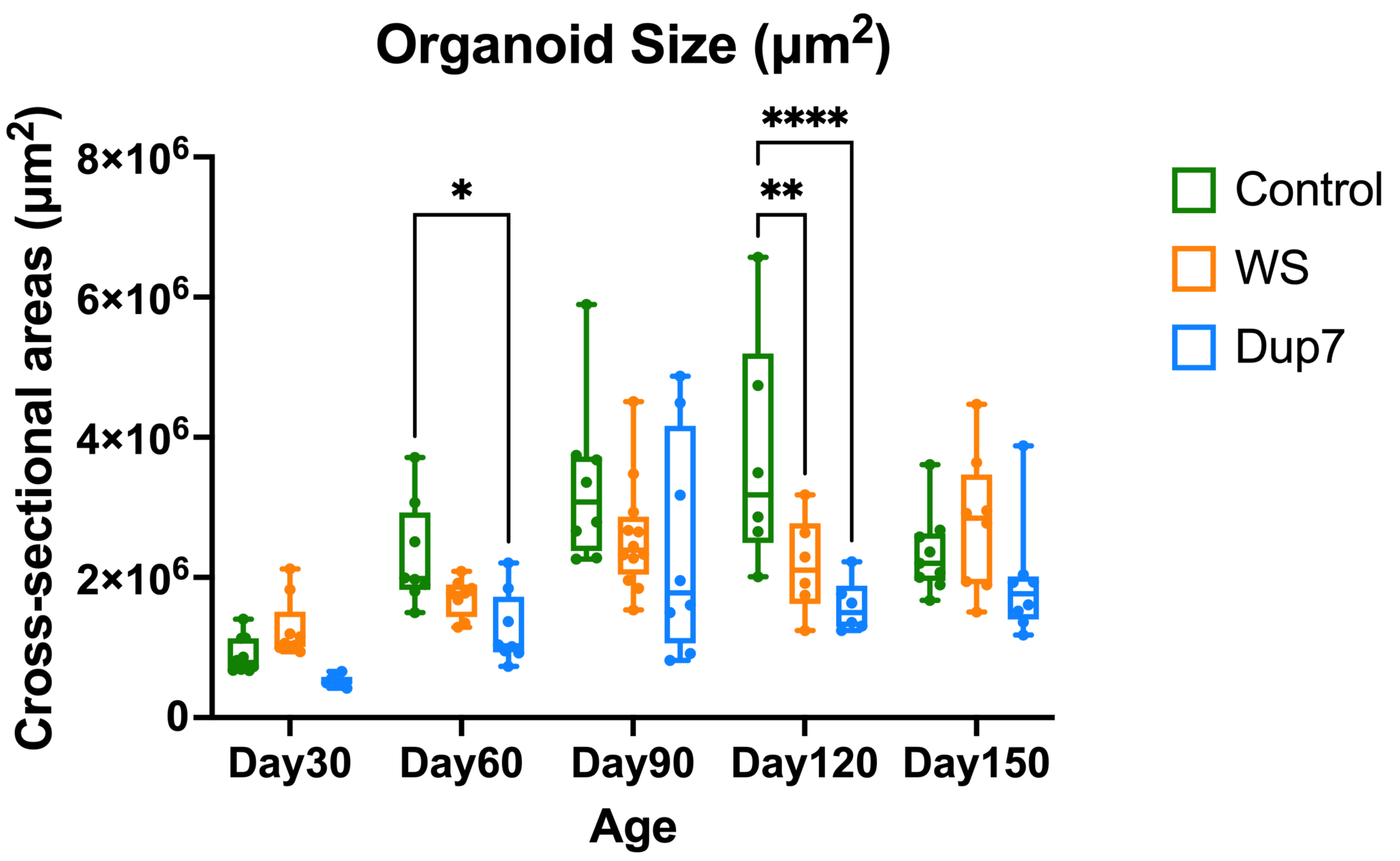


**Supplementary Figure 1. Growth kinetics and area expansion of embryoid bodies.** Longitudinal morphometric analysis of embryoid body (EB) size ($\mu m^2$) from Day 1 to Day 55. Each data point represents the cross-sectional area of an individual EB. Boxes indicate the 25th–75th percentiles, and whiskers represent the minimum to maximum values.

**Supplementary Figure 2. Comparison of organoid cross-sectional area across genotypes and developmental stages.** Longitudinal analysis of organoid size ($\mu m^2$) from Day 30 to Day 150. Each data point represents the cross-sectional area ($\mu m^2$) of an individual organoid. Boxes indicate the 25th–75th percentiles, and whiskers represent the minimum to maximum values. Two-way ANOVA revealed significant main effects for both Age (P-value less than 0.0001) and Genotype (P-value less than 0.0001), along with a significant Interaction effect (P-value = 0.0316). Data represent predicted means (LS means) derived from 120 total values. Statistical significance was evaluated at alpha = 0.05 (* P-value less than 0.05, **** P-value less than 0.0001).

**Supplementary Figure 3**

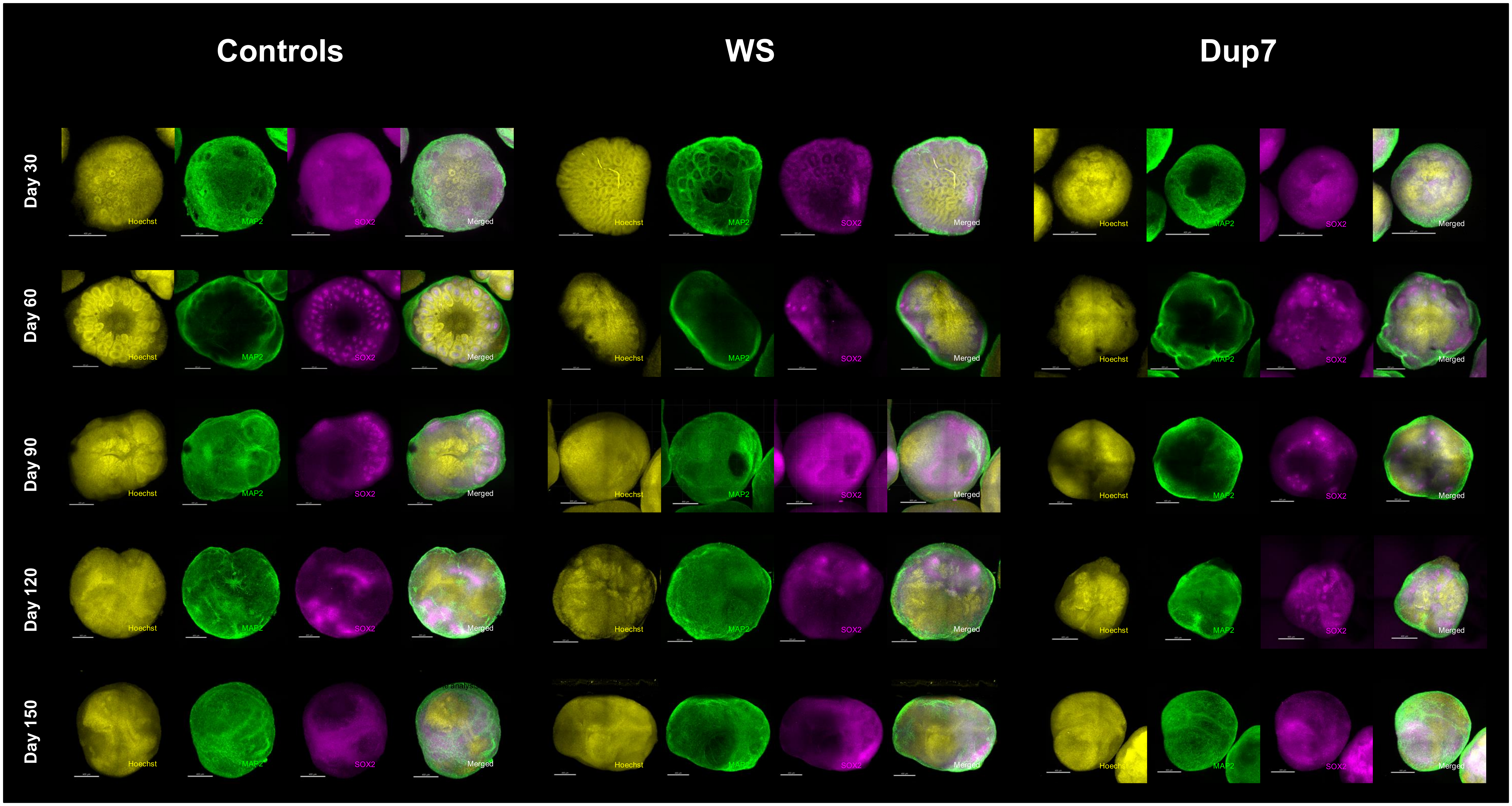


**Supplementary Figure 3. Longitudinal immunohistochemical profiling of cleared organoids.** Representative confocal images of cleared cerebral organoids across Control, WS, and Dup7 genotypes at multiple developmental time points. Tissues were processed through a clearing protocol and immunostained for Hoechst (nuclei; yellow), MAP2 (mature neurons; green), and SOX2 (neural progenitors; magenta). Merged images illustrate the spatial organization of progenitor zones and neuronal layers over time. Scale bars = 400 μm.

Supplementary Figure 4

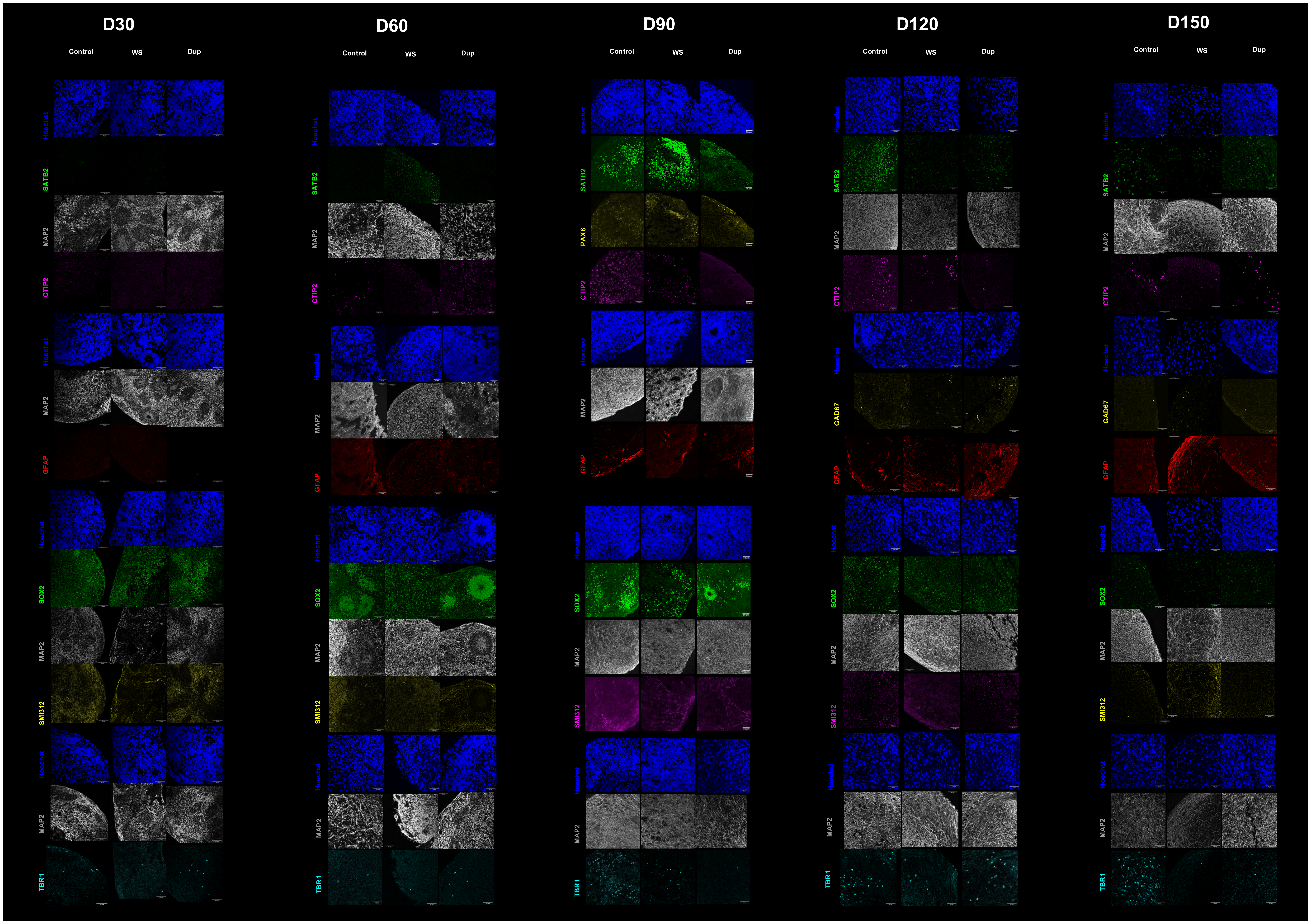


**Supplementary Figure 4. Quality control and lineage-specific immunohistochemical characterization.** Comprehensive immunohistochemical assessment of cell-type diversity and organoid architecture. Representative images of each genotype are shown at various developmental stages to confirm the presence of key neural lineages. Organoids were characterized using the following markers: Progenitors–PAX6 and SOX2; Excitatory Neurons–TBR1 (deep layer), CTIP2 (deep layer), and SATB2 (upper layer); Inhibitory Neurons–GAD67; Axonal Integrity–SMI-312; Astrocytes–GFAP; General Morphology–MAP2 and Hoechst Detailed staining combinations and specific time points are provided in the Methods section. Scale bars = 50 μm.

**Supplementary Figure 5**

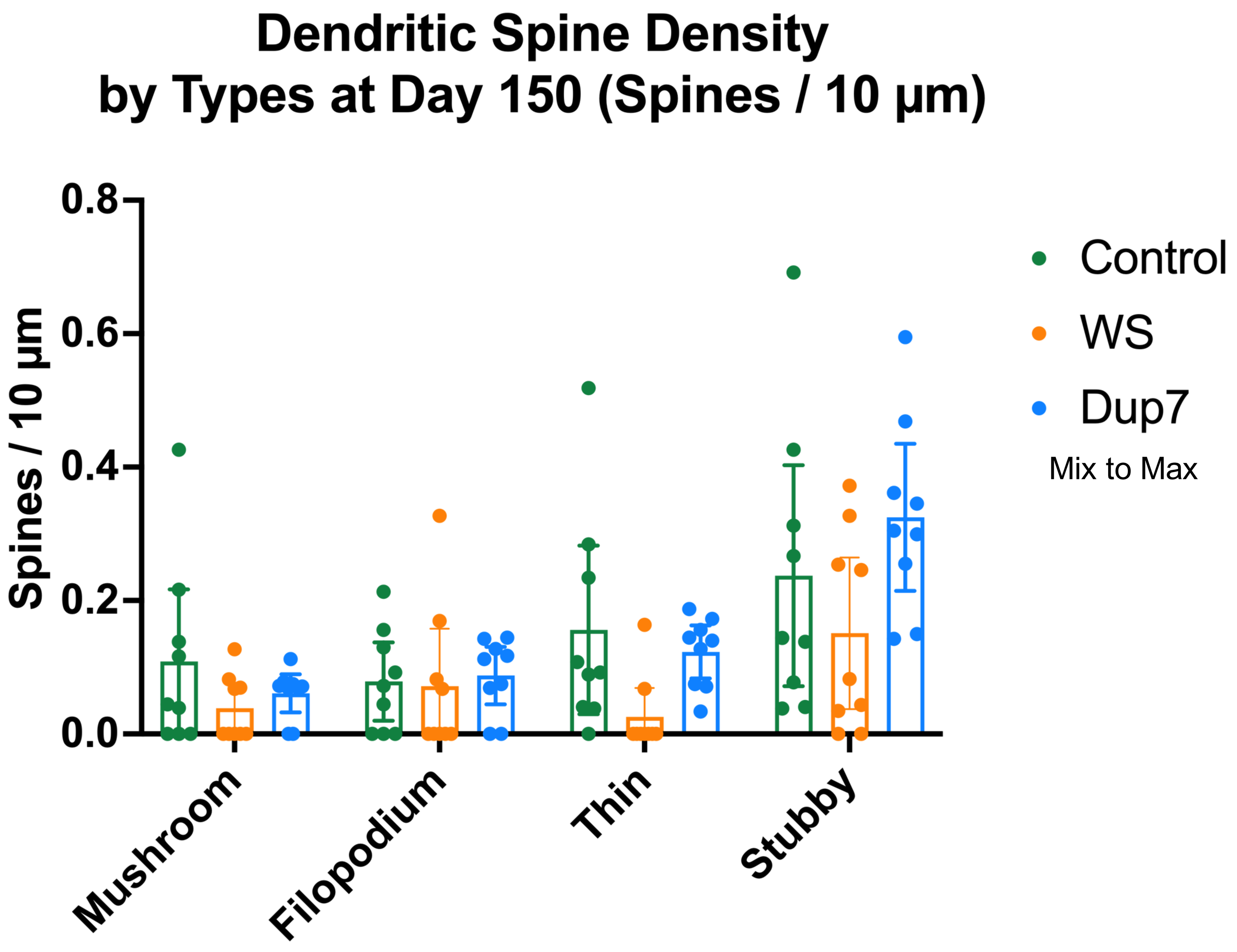


**Supplementary Figure 9. Linear regression analysis of longitudinal gabazine responsiveness across genotypes.** Simple linear regression models and calculated goodness-of-fit parameters for Control, WS, and Dup7 groups across developmental time points. The Control group exhibits an $R^2$ value of 0.5649 (F = 18.17, P = 0.0008) with a calculated X-intercept of 137.7 (95% CI: 118.2 to 184.4). The WS group exhibits an $R^2$ value of 0.2523 (F = 4.723, P = 0.0474) with a calculated X-intercept of 157.1 (95% CI: 123.3 to 3940). The Dup7 group is defined by a vertical line fixed at day 60.

**Supplementary Figure 6**

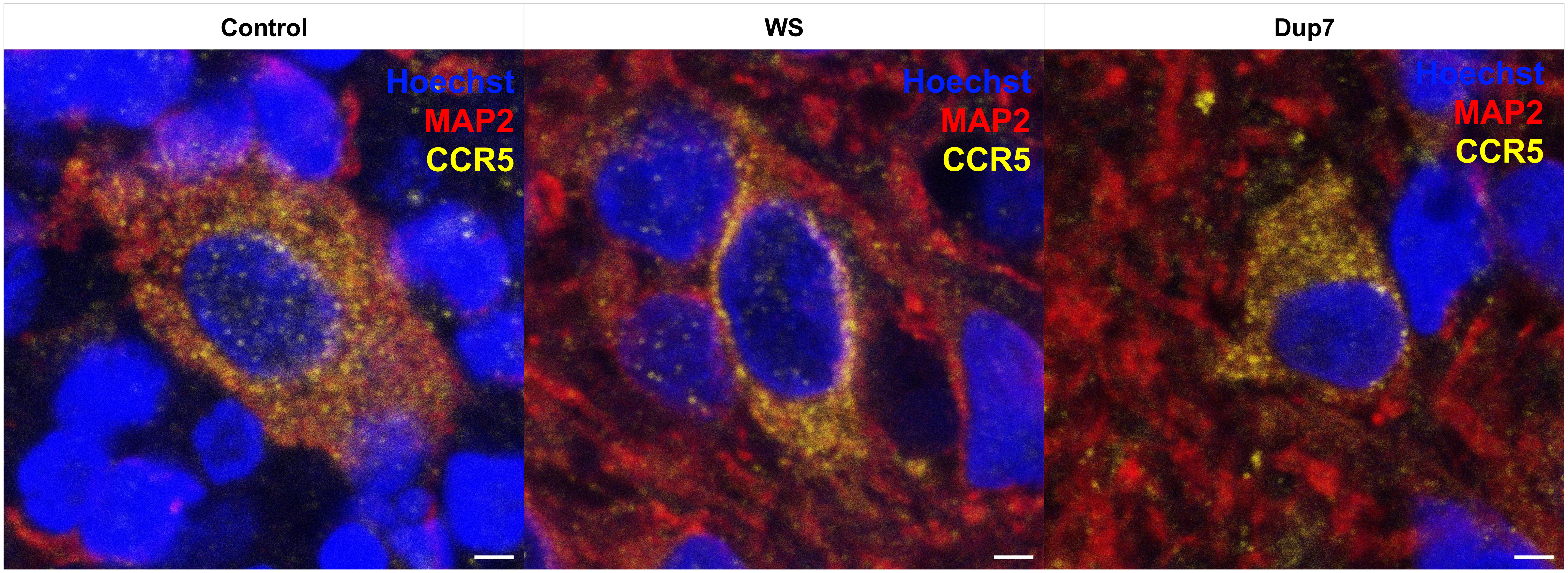


**Supplementary Figure 6. Expression of CCR5 Across Genotypes at Day 150.** Representative immunohistochemical images demonstrating the expression of CCR5 (Yellow) on Day 150 Control, WS, and Dup7 cerebral organoids. Neurons are stained with MAP2 (Red). Nuclei are counterstained with Hoechst (blue). Scale bars = 2 μm.

**Supplementary Figure 7**

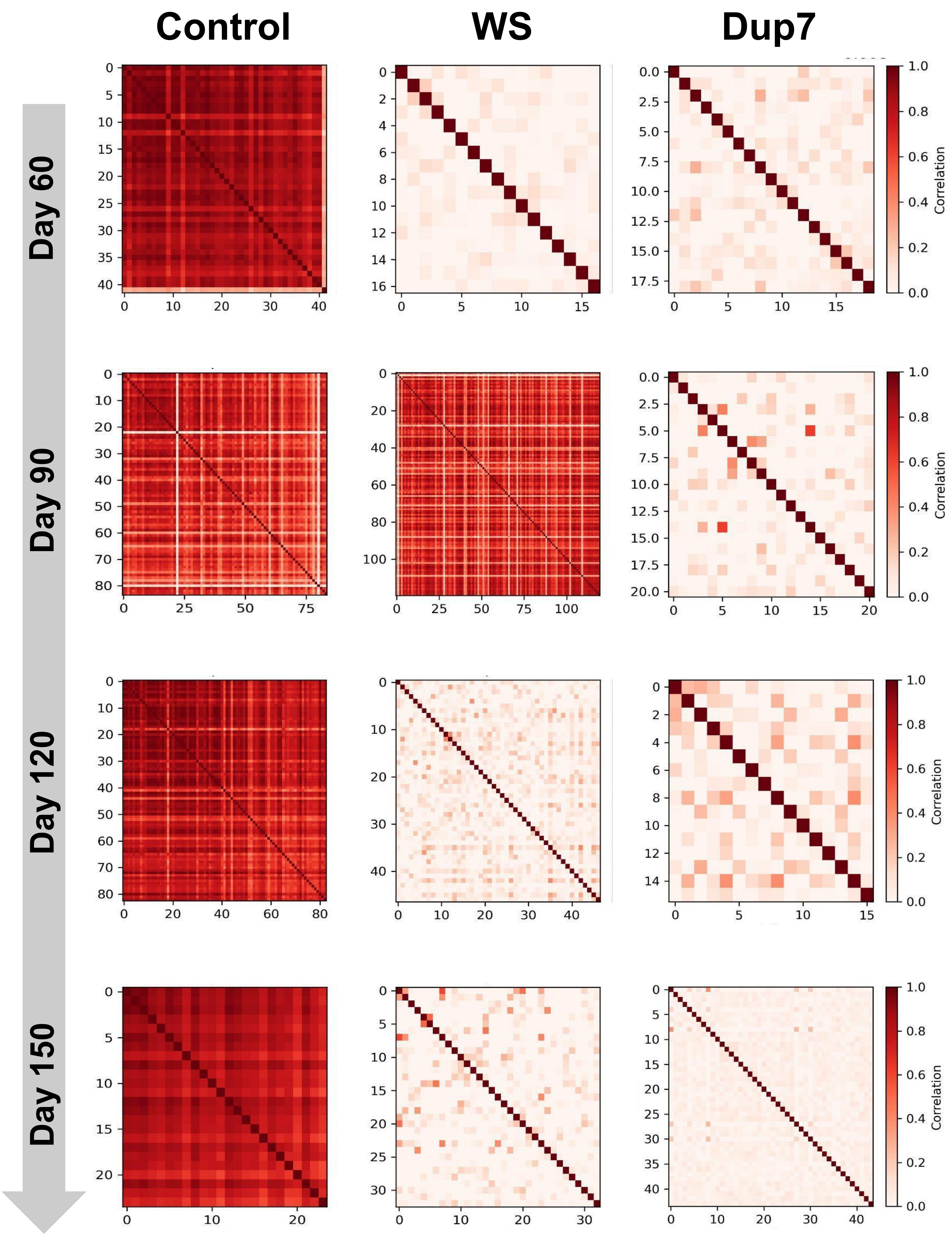


**Supplementary Figure 7.** Representative Cross-Correlation plots of spontaneous functional activity across three genotypes (Control, WS, and Dup7) at Day 60, 90, 120, and 150.

**Supplementary Figure 8**

### Matched Pre/Post Gabazine Analysis in Neuronal Activity and Functional Cross-Correlation

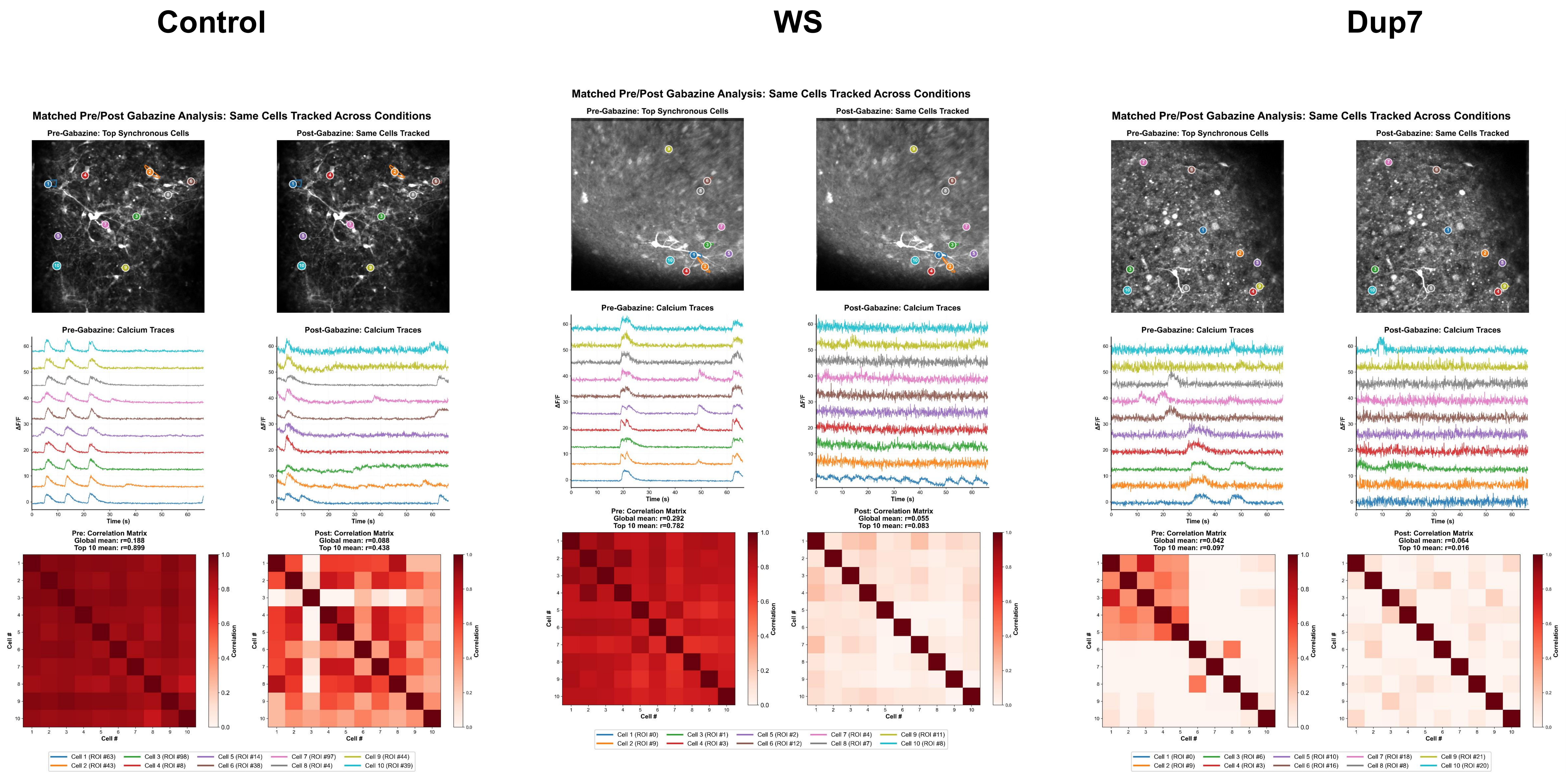


**Supplementary Figure 8. Functional network alterations and Gabazine-mediated rescue in 7q11.23 CNV organoids.** Single-cell functional dynamics and GABAergic response. Top row: representative average projection images of individual images and ten identified neurons; second row: corresponding calcium transients; bottom row: functional cross-correlation heatmaps. Results are shown for pre- and post-Gabazine treatment within each genotype at Day 60.
